# Members of the CUGBP Elav-Like Family of RNA-Binding Proteins are Expressed in Distinct Populations of Primary Sensory Neurons

**DOI:** 10.1101/2021.07.05.451166

**Authors:** Eliza Grlickova-Duzevik, Thomas M Reimonn, Merilla Michael, Tina Tian, Jordan Owyoung, Aidan McGrath-Conwell, Peter Neufeld, Madison Mueth, Derek C Molliver, Patricia Jillian Ward, Benjamin J Harrison

## Abstract

Primary sensory Dorsal Root Ganglia (DRG) neurons are diverse, with distinct populations that respond to specific stimuli. Previously, we observed that functionally distinct populations of DRG neurons express mRNA transcript variants with different 3’ untranslated regions (3’UTR’s). 3’UTRs harbor binding sites for interaction with RNA-binding proteins (RBPs) for transporting mRNAs to subcellular domains, modulating transcript stability and regulating the rate of translation. In the current study, analysis of publicly available single-cell RNA-Sequencing (scRNA-Seq) data generated from adult mice revealed that 17 3’UTR-binding RBPs were enriched in specific populations of DRG neurons. This included 4 members of the CUGBP Elav-Like Family (CELF): CELF2 and CELF4 were enriched in peptidergic, CELF6 in both peptidergic and nonpeptidergic and CELF3 in tyrosine hydroxylase-expressing neurons.

Immunofluorescence studies confirmed that 60% of CELF4+ neurons are small diameter C fibers and 33% medium diameter myelinated (likely Aδ) fibers and showed that CELF4 is distributed to peripheral termini. Co-expression analyses using transcriptomic data and immunofluorescence revealed that CELF4 is enriched in nociceptive neurons that express GFRA3, CGRP and the capsaicin receptor TRPV1. Re-analysis of published transcriptomic data from macaque DRG revealed a highly similar distribution of CELF members, and re-analysis of single-nucleus RNA sequencing data derived mouse and rat DRG after sciatic injury revealed differential expression of CELFs in specific populations of sensory neurons. We propose that CELF RNA-binding proteins may regulate the fate of mRNAs in populations of nociceptors, and may play a role in pain and/or neuronal regeneration following nerve injury.

## INTRODUCTION

RNA binding proteins (RBPs) are critical for orchestrating the post transcriptional fate of transcripts, including transport to sub-cellular domains for local translation, storage and release from ribonucleoprotein complexes and recruitment of translational machinery (e.g., review: (Lenzken et al., 2014)). The 3’ untranslated region (3’UTR) of transcripts is a hub for mRNA-protein interactions (e.g., review:(Szostak & Gebauer, 2013)). For example, RBP binding to 3’UTRs on nascent transcripts regulates poly(A) signal site selection, cleavage and polyadenylation (e.g., review: (Tian & Manley, 2013). On mature transcripts, 3’UTRs contain interaction sites (originally termed “zip-code” motifs (Kislauskis et al., 1994; Kislauskis & Singer, 1992; Ross et al., 1997), for subcellular trafficking (Arora et al., 2022; Kim et al., 2015; Ribeiro et al., 2020; Yergert et al., 2021) anchoring points for scaffolding interactions (Sahoo et al., 2018) and interaction sites for recruitment of miRNAs (Grimson et al., 2007; Majoros & Ohler, 2007) or inhibition by competition for binding sites (Srikantan et al., 2012).

3’UTR-protein interactions are highly dynamic and provide a regulatory layer for modulation of diverse cellular functions. By interaction with alternative cleavage sites at the distal end of nascent transcripts, RBP-3’UTR interactions generate alternate 3’UTR isoforms through a mechanism termed Alternative Polyadenylation (APA) (Singh et al., 2009). mRNA isoforms with long 3’UTRs are predominantly expressed during development (Ji & Tian, 2009), and in the adult are restricted to certain cell types most notably neurons that express mRNAs with long 3’UTRs (Harrison et al., 2014). Differential expression of 3’UTR isoforms by APA is associated with diverse biological functions, for example long 3’UTR isoforms upregulated during neuroplasticity (An et al., 2008; Miura et al., 2013) and during responses to stress (Graber et al., 2013; Zheng et al., 2018), etc. 3’UTR variants are associated with human disease (Griesemer et al., 2021). For example, shortened 3’UTR sequences contain fewer interactions sites for micro-RNAs, and are associated with neurological disease such as Huntington’s (Romo et al., 2017), and facilitate oncogene activation and tumor growth (Mayr & Bartel, 2009).

RBP-3’UTR interactions are especially important for remote cellular compartments that rely on a supply of mRNA for local protein synthesis. Sensory neurons have extremely long axons projecting from peripheral targets through their cell bodies (soma) housed in DRG terminating in the spinal cord dorsal horn. These termini require a supply of raw materials and machinery that is transported along neuronal processes. RBPs are required for mRNA transport to and translation in these remote domains for maintenance of synapses and for plasticity (Hornberg & Holt, 2013; Wagnon et al., 2012). Numerous “neuron-specific” RBPs have been identified that are highly enriched in neurons and are required for mRNA transport and neuronal excitability (Darnell, 2013; Hornberg & Holt, 2013).

Primary somatosensory neurons are classified according to their responsiveness to specific stimuli (light touch, temperature, stretch, etc.), neurochemistry and neurophysiology (Le Pichon & Chesler, 2014), and can be identified by population-specific histological markers. Sensory neurons with unmyelinated axons (C-fibers) tend to have small-diameter cell bodies, and the great majority of these neurons are considered to be nociceptors (reviewed (Light, A.R. 1992)), grossly divided into two subpopulations: peptidergic neurons, which express Calcitonin Gene Related Peptide (CGRP) and Ntrk1 (gene for TRKA) and are particularly important for the transduction of noxious heat stimuli, and non-peptidergic neurons, which bind the plant lectin IB4 and express the G protein-coupled receptor MAS Related GPR Family Member D (MRGPRD) and are particularly important for mechanical nociception (Dong et al., 2001; Silverman & Kruger, 1988). Most sensory neurons that express the heat- and acid-gated channel TRPV1 in adulthood are peptidergic neurons (Zwick et al., 2002). Peptidergic neurons also include a subset of thinly myelinated (A-delta) nociceptors and a minority of A-beta afferents (Lawson et al., 1993, 1996). A third population of C-fibers is comprised of neurons that express tyrosine hydroxylase (TH) and are not nociceptors; instead they are sensitive mechanoreceptors that have been implicated in the transduction of pleasant “social touch” (Lallemend & Ernfors, 2012; Li et al., 2011). Although markers of the peptidergic/non-peptidergic nociceptor populations are highly segregated in the mouse, the selectivity of these markers varies considerably across species; in mouse, IB4-binding is seen in less than 10% of neurons expressing CGRP or TRPV1, whereas in the rat this overlap is 35% (Price & Flores, 2007; Zwick et al., 2002), and recent evidence shows that in humans CGRP and TRPV1 are expressed in most nociceptors (IB4 does not bind sensory neurons in human) (Shiers et al., 2021). Recent advances in single-cell RNA-sequencing (scRNA-Seq) allow sensory neurons to be characterized by comprehensive mRNA expression profiles, rather than a handful of neurochemical markers. These transcriptomic studies align closely with histological data and are a powerful resource for identifying the genomic signatures of functionally-distinct neuronal populations (Usoskin et al., 2015).

Previously, we reported that functionally distinct populations of DRG neurons express mRNA isoforms with divergent 3’UTR sequences (3’UTR isoforms), and that these 3’UTR variants contain different RBP interaction motifs (Harrison et al., 2019). This led us to theorize that sensory neuron diversity is controlled by differential RBP-3’UTR interactions. In the current study we examine the expression of 3’UTR-binding RBPs in DRG, providing further evidence suggesting divergence of 3’UTR-protein interactions in distinct populations of sensory neurons.

## METHODS

### Animals

All animal procedures were approved by the Institutional Animal Care and Use Committee of the University of New England, and Emory University consistent with federal regulations and guidelines. All animals used for this study were 6-8 week old male and female C57/BL6 mice purchased from Jackson Labs (Bar Harbor, Maine).

### Dorsal Root Ganglion Single Cell RNA-seq Analysis

#### Trimming, aligning, and quantification

Previously published single-cell RNA-seq data from adult mouse dorsal root ganglion neurons produced by Usoskin et al. was accessed from GEO using accession code GSE59739 (Usoskin et al., 2015). Fastq files were downloaded and reads were trimmed using Trimmomatic version 0.39 (Bolger et al., 2014) in single-end mode with the following parameters: ILLUMINACLIP:TruSeq3-SE:2:30:10 LEADING:3 TRAILING:3 SLIDINGWINDOW:4:15 MINLEN:36. Trimmed reads were aligned to the mouse genome (GRCm38, primary assembly) and transcripts mapping to genes were quantified (Gencode vM25, primary assembly) using STAR version 2.7.5a (Dobin et al., 2013) with default parameters. Transcript quantifications were summed by gene when one cell contained reads from multiple fastq files, and gene counts were tabulated into a gene-by-cell expression matrix.

#### Clustering, Visualization, and Annotation

Downstream processing was performed using Seurat version 3.2.0 (Stuart et al., 2019). Cells that expressed fewer than 500 genes, expressed more than 10,000 genes, contained fewer than 1,000 transcripts, or had more than 10% of transcripts mapping to mitochondrial genes were removed, resulting in 811 cells passing all metrics. Gene counts were normalized using “LogNormalize” and counts per 10,000 normalization. 3,000 highly variable genes were selected using the “vst” method. Normalized transcript counts were scaled to 0 mean and unit variance, and principal component analysis with 100 principal components was performed. Cells were Louvain clustered with a resolution of 1 and k = 20 nearest neighbors resulting in six clusters. Cells were embedded using t-SNE and UMAP with default parameters. A grid search of clustering parameters for k nearest neighbors in {10, 15, 20, 30} and resolution in {0.5, 1, 1.5, and 2} was performed with visual inspection of concordance between UMAP embedding and clustering to verify that k = 20 and resolution = 1 were reasonable. Normalized gene expression for 12 marker genes (Th, Sst, Mrgprd, P2rx3, Tac1, Ntrk1, Calca, Nefh, Pvalb, B2m, Vim, Col6a2) were plotted and cluster cell types were manually annotated as Non-neuronal, Peptidergic (Pep), non-peptidergic (NP), tyrosine hydroxylase-expressing (TH) and neurofilament heavy chain (Nefh)-expressing (NF) (Fig 1). To identify cell subpopulations, each cell type was separately processed using 2000 variable features, 30 principal components, k = 10 nearest neighbors, and a louvain resolution of 1.

**Figure 1:**
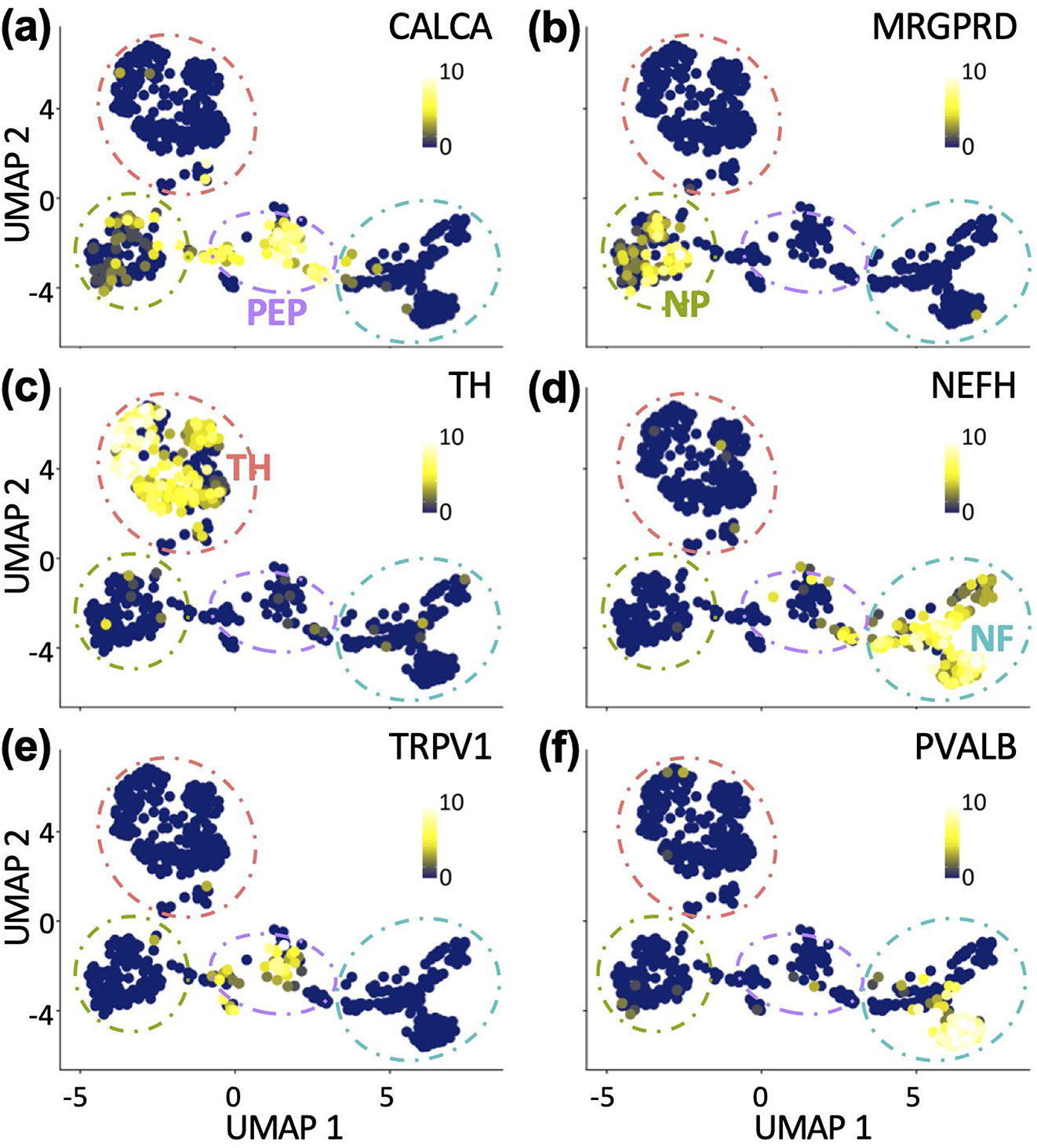
Utilization of publicly available single cell RNA sequencing (scRNA-Seq) data, derived from adult mouse sensory neurons, to generate population-specific gene expression profiles. To generate gene expression profiles for specific populations of mouse adult sensory neurons, publicly available scRNA-seq data (GSE59739) were downloaded and processed with the latest release of SEURAT software using up to date gene annotations. Graphs are Uniform Manifold Approximation and Projections (UMAPs) of gene expression profiles. Each data point is a cell in UMAP space. Yellow shading = high expression, Blue shading = low expression. Expression profiles distributed into clusters as expected: **A-D** Peptidergic neurons were identified by CALCA (CGRP) expression (PEP, purple cluster), non-peptidergic by MRGPRD expression (NP, green cluster), tyrosine hydroxylase–expressing neurons (TH, red cluster) and myelinated neurons by neurofilament heavy chain expression (NF, blue cluster). **E** Sub-cluster of TRPV1-expressing nociceptors. **F** Sub-cluster of parvalbumin (PVALB) -expressing proprioceptors.

#### Differential Expression Analysis

Differentially expressed genes were identified using a Wilcoxon rank sum test for each cell type compared to all other cell types where the log2 fold change was greater than 0.25 and the gene was expressed in at least 10% of cells. A list of all known 3-UTR-binding proteins was obtained from the Gene Ontology consortium website (Ashburner et al., 2000; Gene Ontology Consortium, 2021) to highlight differentially expressed cell type specific 3’UTR-binding proteins.

#### mRNA co-expression analysis

Pearson’s correlation was calculated by comparison of CELF4 mRNA expression values vs expression values for all other detected mRNAs. For reference, also shown are values for CGRP vs NTRK1 (known to be co-expressed in the same population of neurons) and CGRP vs MRGPRD (known to be expressed separately in distinct populations) (Fig 6A).

### Antibody characterization (Table 1)

Polyclonal rabbit anti-CELF4 antibody (Prestige Antibodies, Sigma-Aldrich, HPA037986): Developed using CUGBP, Elav-like family member 4 recombinant protein epitope signature tag (PrEST). This antibody has previously been extensively validated for use with immunostaining: This antibody detected CELF4 protein in the wildtype mouse hippocampal and cortical neurons, but not in the CELF4 knockout samples (Wagnon et al., 2011, 2012). In addition, we determined that CELF4 gene deletion (CELF4-fl/fl (Wagnon et al., 2011) ; Avil-cre-ERT2 mice treated with tamoxifen (Lau et al., 2011) resulted in loss of signals (Fig 4A), as did omission of primary antibody (Fig 6C, 7), providing further evidence that this reagent is specific and robust.

**Table 1:** Primary antisera used for this study.

| *Target* | *Immunogen* | *RRID* | *Host* | *Supplier* | *Cat #* | *Dilution Factor* |
| --- | --- | --- | --- | --- | --- | --- |
| CELF4 | MYIKMATLANGQA<br>DNASLSTNGLGSSPG<br>SAGHMNGLSHSPG<br>NPSTIPMKDH | AB_10673035 | Rabbit | Millipore Sigma | HPA037986 | 1:800 |
| TRPV1 | YTGSLKPEDAEVFKD<br>SMVPGEK | AB_1624142 | Guinea Pig | Neuromics | GP14100 | 1:500 |
| NF-H | NF-H purified from<br>bovine spinal cord | AB_2149761 | Chicken | EnCor | CPCA-NF-H | 1:40000 |
| CGRP | ACDTATCVTHRLAGL<br>LSRSGGVVKNNFVPT<br>NVGSKAF | AB_1282813 | Guinea Pig | Fitzgerald Industries<br>International | 20R-CP007 | 1:1500 |

Polyclonal guinea pig anti-TRPV1 (Neuromics, GP14100): The antigen was residues of the carboxy-terminus of rat TRPV1, YTGSLKPEDAEVFKDSMVPGEK. This antibody has been assessed with wild-type tissue, as well as TRPV1-deficient mice, and no apparent immunoreactivity was observed in TRPV1-KO mouse DRG, although the antibody robustly stained the expected population of wild-type neurons (Sand et al., 2015). In addition, omission of primary antibody resulted in loss of signals (Fig 6C).

Polyclonal chicken anti NF-H (EnCor, CPCA-NF-H): NF-H antibody robustly stained the expected population of medium and large diameter DRG neurons and large caliber axons. No signal was detected in NF-H negative small diameter neurons, small caliber fibers or non-neuronal cells (Fig 5), thus confirming the specificity of this reagent.

Polyclonal guinea pig anti-CGRP (20R-CP001): CGRP antibody was raised in guinea pig using calcitonin gene-related peptide conjugated to BSA as the immunogen. This antibody has been extensively used for immunostaining DRG tissue slices to identify peptidergic neurons, and specifically stains this population (Harrison et al., 2014).

### Immunofluorescence Imaging and Analysis

Immunofluorescent staining was used to characterize populations of DRG neurons that express CELF4 using reagents detailed in Table 1: CGRP for peptidergic, TRPV1 for heat responsive nociceptors, and NF-H for myelinated neurons. Additionally, the plant lectin IB4 conjugated to fluorophore was used to stain the non-peptidergic population, e.g.: (Burnstock, 2000)

#### Sample Collection and Processing

Animals were exsanguinated under deep pentobarbital anesthesia, before perfusion with cold 4% paraformaldehyde in PBS. DRG were immediately dissected, fixed in 4% paraformaldehyde (PFA) in PBS for 10 minutes before cryoprotection by incubating overnight in 30% sucrose. 15 μm sections were cut using a cryostat, mounted onto adhesion microscope slides, and left to dry, uncovered at room temperature for at least 1 hour.

#### Immunostaining

Non-specific binding sites were blocked by incubating sections with 5% donkey serum in 0.4% Triton X-100 PBS for 1 hour, covered. CELF4 antibody was diluted 1:800, TRPV1 1:500, CGRP 1:1000 and/or NF-H 1:40000 in blocking buffer and left on sections overnight (i.e. 18-24 hours) at room temperature. The sections were then washed with 0.4% Triton X-100 PBS; the washing solution was left on for 5 minutes for the first wash, then 10 minutes for the following three washes. Secondary antibodies and counterstains were prepared 1:200 in blocking buffer, added to sections and for 2 hours at room temperature. Alexa 488-conjugated IB4 was diluted at 1:50. The sections were washed a second time using the same procedure as the first wash. Slides were coverslipped with an antifade mountant and left overnight at room temperature before imaging.

#### Imaging

Multichannel fluorescent micrographs of entire longitudinal DRG sections were obtained using a Leica DMi8 epifluorescent inverted microscope with 20X objective and automated field stitching. Image analysis was performed as previously published (Harrison et al., 2014): Two sections per animal containing more than 200 neurons with visible nuclei were selected from four animals. Sections were spaced more than 50 μm apart to ensure that no single neuron could be included in the analysis twice. Using Image J software (Schneider et al., 2012), individual neurons with visible nuclei were then manually circled to generate separate regions of interest (ROIs). Mean fluorescent intensities were recorded along with the cross-sectional area from each ROI. All ROI data from both sections from individual animals were then collated into a single dataset per animal, yielding data from over 400 neurons per animal. Background was estimated as the mean signal intensity of non-expressing neurons. The median staining intensity for each channel was then calculated for each of the four datasets (i.e., the median intensity of each DRG) to allow for normalization of fluorescence intensities to the mean median value (division by central tendency [median] method). First, to observe the distribution of CELF4 expression according to neuron soma size, normalized signals from all animals were collated into a single dataset and subsequently binned according to soma cross-sectional area (50μm bin sizes) (Fig. 4C). Neurons were assigned to either small (<300μm), medium (300-700μm) or large (>700μm) populations. These size divisions were chosen due to the well-established somatosensory characteristics that vary according to soma size. To identify peptidergic, non-peptidergic and myelinated populations, sections were co-stained with CGRP antibody, IB4 lectin or neurofilament heavy chain (NF-H) antibody respectively (Ruscheweyh et al., 2007).

#### Neuronal Retrograde Tracing

For retrograde labeling of muscle afferents, 8 adult wild type C57BL/6J mice (Jackson Laboratories, strain #000664) at least 8-weeks of age were studied. The right bilateral heads of the gastrocnemius and tibialis anterior muscles received 2μL injections of 5.0% fast blue retrograde tracer (Polysciences, 17740) with a 26-gauge 10μL Hamilton microliter syringe (Model #701) distributed in at least 3 areas within the muscle bellies under 2% isoflurane (Piramal Enterprises Limited, Telangana, India, ANADA 200-237, NDC 66794-013-25) in 1L/min O_2_, totaling 6μL of tracer. All mice received 1 dose of 5mg/kg 1.5mg/ml meloxicam (Pivetal, ANADA 200-550, NDC 46066-170-13). Four to 14 days later, to allow for retrograde transport, mice were euthanized and perfused with 0.4% paraformaldehyde as described above. The right L4 dorsal root ganglion (DRG) was dissected and placed in 20% sucrose for cryoprotection prior to being cryosectioned for immunohistochemistry as detailed above.

#### Skin Imaging

For analysis of lanceolate endings, 5 additional wild type C57BL/6J mice were used. Following euthanasia and paraformaldehyde perfusion, a quadrangular piece of hairy hind paw skin was dissected, cryoprotected and sectioned transversely at 30 μm. Following immunolabeling with NF-H and CELF4, the skin was imaged on a Nikon A1R with 60x objective, 0.5 μm step size.

## RESULTS

To identify RBPs expressed in populations of sensory neurons, we obtained publicly available single-cell RNA-Sequencing data generated from adult mouse DRG (GEO GSE59739) (Usoskin et al., 2015). Since release of those data, analysis tools have evolved significantly. In addition, gene annotations, mapping coordinates and transcript variant sequences are under constant revision. We therefore reanalyzed the raw sequencing data using current software (SEURAT 3.2.0) and annotations (Gencode vM25). Cells clustered into functionally defined populations as expected (Fig 1): Peptidergic (PEP) that are predominantly nociceptors that express neuropeptides such as CGRP, non-peptidergic (NP) that are predominantly nociceptors that do not express these neuropeptides, tyrosine hydroxylase-expressing (TH) small-diameter sensory neurons implicated in the pleasurable, affective component of touch and neurofilament heavy chain (NF-H)-expressing that are myelinated and respond to low threshold stimuli such as light touch. Within those clusters, neurons grouped into expected sub populations, for example TRPV1-expressing neurons (heat responsive nociceptors) subclustered within the PEP cluster, and parvalbumin-expressing neurons (chiefly proprioceptors) subclustered within the NF (myelinated) cluster (Fig 1) (Usoskin et al., 2015).

Gene expression profiles from each population (Pep, NP, TH, NF) were compared by differential gene expression analysis using Wilcoxon rank sum test with SEURAT software. Of approximately 1100 known RBPs annotated in the mouse genome, 100 are assigned the gene ontology (GO) term “mRNA 3’-UTR binding” (Table 2). Of those, 84 3’UTR-binding proteins were detectable in DRG neurons (expression > 1 CFM) and 17 were enriched in at least 1 population (Table 2). Therefore, the majority (80%) of 3’UTR-binding proteins detected in DRG neurons were ubiquitously expressed throughout all populations, as exemplified by FXR1 (Fig 2 a,b). Interestingly, 4 members of the CUGBP Elav-Like Family (CELF) were differentially expressed: CELFs 2, 3, 4 and 6 (Table 3, Fig 2). CELF2 was enriched in the PEP population, CELF3 in the TH population, CELF4 in the PEP population and CELF6 was enriched in both NP and PEP populations.

**Table 2:** Function of 3’UTR mRNA-binding proteins in populations of adult mouse sensory (DRG) neurons. 100 mouse genes are assigned the “3’UTR-binding” ontology term (Ashburner et al., 2000). mRNA’s for 84 of these were detected in publicly available scRNA-Seq data derived from adult mouse DRG neurons (GSE59739, (Usoskin et al., 2015). 17 were significantly differentially expressed (FDR<0.05) between one or more of 4 DRG neuron populations (tyrosine hydroxylase-expressing, peptidergic, non-peptidergic, myelinated).

| *Gene Ontology (GO) Term* | *3'UTR-binding proteins (100)* | *Detected in DRG neurons (84)* | *Enriched in DRG neuron population(s) (17)* |
| --- | --- | --- | --- |
| posttranscriptional regulation of gene expression (GO:0010608) | 8 | 8 | 1 |
| regulation of mRNA metabolic process (GO:1903311) | 33 | 32 | 8 |
| regulation of translation (GO:0006417) | 43 | 35 | 4 |
| regulation of mRNA catabolic process (GO:0061013) | 33 | 11 |  |
| regulation of mRNA stability (GO:0043488) | 39 | 33 | 5 |
| mRNA processing (GO:0006397) | 45 | 43 | 10 |
| RNA splicing (GO:0008380) | 32 | 31 | 10 |
| RNA localization (GO:0006403) | 5 | 4 |  |
| regulation of gene silencing by miRNA (GO:0060964) | 10 | 9 |  |
| RNA transport (GO:0050658) | 10 | 7 | 1 |
| cytoplasmic ribonucleoprotein granule (GO:0036464) | 34 | 28 | 4 |
| growth cone (GO:0030426) | 6 | 6 | 2 |
| dendritic spine (GO:0043197) | 5 | 5 | 1 |
| ribosome (GO:0005840) | 6 | 6 | 2 |
| distal axon (GO:0150034) | 4 | 4 |  |
| nucleolus (GO:0005730) | 16 | 16 | 1 |
| dendrite (GO:0030425) | 13 | 12 | 2 |
| postsynaptic density (GO:0014069) | 7 | 7 | 1 |
| nuclear body (GO:0016604) | 13 | 13 | 2 |
| neuronal cell body (GO:0043025) | 10 | 10 | 3 |
| postsynapse (GO:0098794) | 8 | 8 | 2 |

**Table 3:** 3’UTR mRNA-binding proteins significantly enriched (FDR<0.05) in DRG neuron populations, sorted by average fold enrichment. TH = tyrosine hydroxylase-expressing, NP= non-peptidergic, NF= neurofilament heavy chain-expressing and PEP= peptidergic neurons.

| *Gene* | *Population* | *Average*<br>*Enrichment* | *p* |
| --- | --- | --- | --- |
| CELF4 | PEP | 28.11 | 4.07E-12 |
| CARHSP1 | NP | 15.76 | 8.95E-63 |
| RBMS3 | TH | 5.52 | 4.68E-33 |
| AUH | NF | 3.64 | 1.67E-35 |
| CELF6 | PEP | 2.98 | 2.25E-06 |
| CELF6 | NP | 2.94 | 1.86E-20 |
| CELF2 | PEP | 2.58 | 0.002704 |
| CELF3 | TH | 2.53 | 1.35E-21 |
| HNRNPA2B1 | TH | 2.21 | 1.65E-10 |
| FXR2 | NF | 2.02 | 1.24E-14 |
| ELAVL4 | TH | 2.02 | 5.72E-13 |
| CSDC2 | NF | 1.93 | 1.37E-25 |
| FXR2 | NP | 1.92 | 2.30E-05 |
| RBMS1 | NP | 1.88 | 2.24E-13 |
| RBFOX1 | TH | 1.78 | 0.000245 |
| FXR2 | PEP | 0.56 | 0.006902 |
| AUH | NP | 0.46 | 0.003118 |
| YBX3 | NF | 0.45 | 0.007494 |
| TAF15 | NF | 0.40 | 2.32E-15 |
| CELF6 | NF | 0.36 | 1.05E-07 |
| CELF4 | TH | 0.31 | 1.38E-05 |
| RBMS3 | NP | 0.29 | 3.77E-05 |
| HNRNPL | NF | 0.23 | 4.45E-05 |
| RPS7 | NF | 0.20 | 0.000137 |
| CARHSP1 | NF | 0.18 | 0.000383 |
| CARHSP1 | TH | 0.17 | 3.98E-09 |

**Figure 2:**
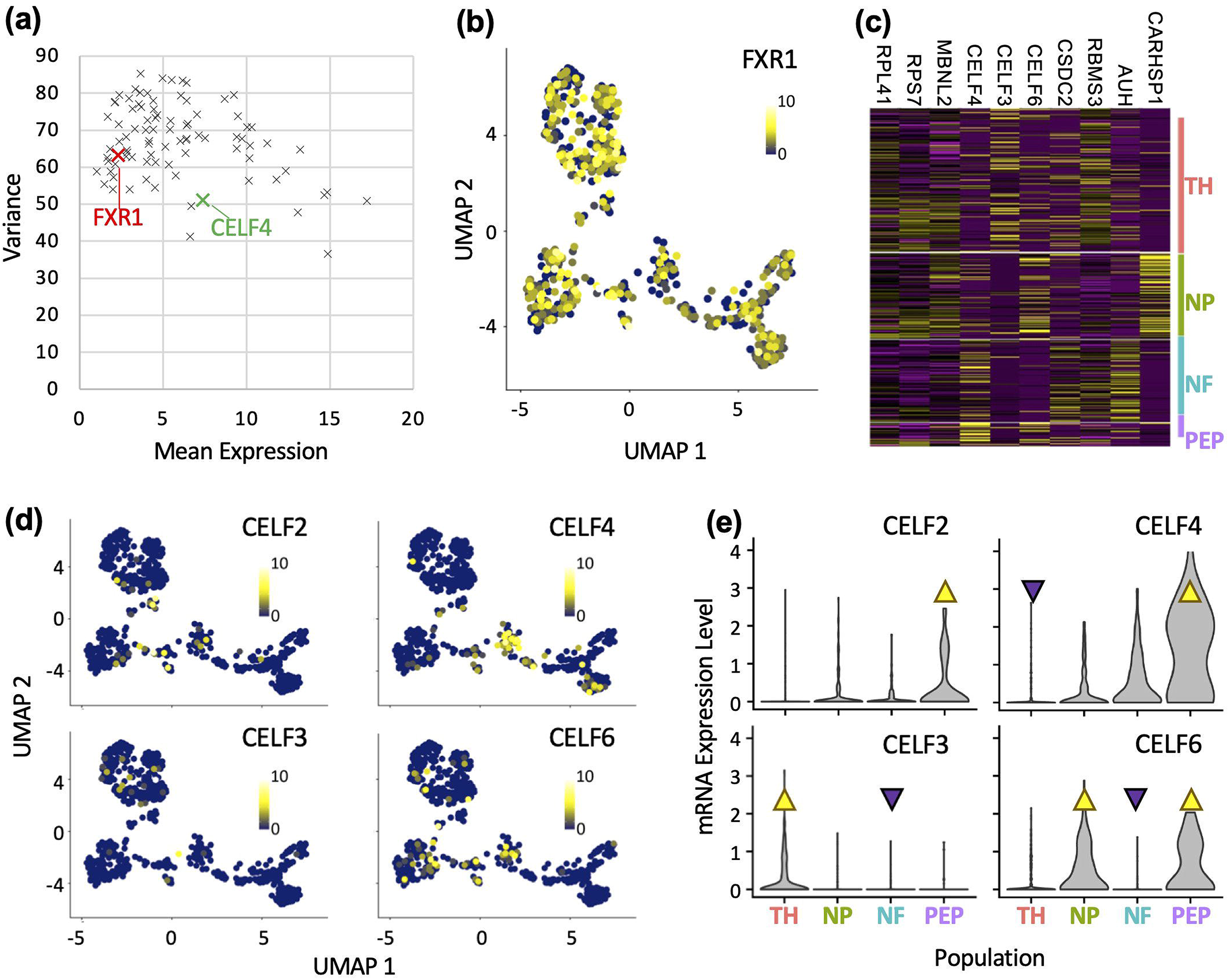
The CELF family of RNA-binding proteins are expressed in distinct populations of sensory neurons. **A** Mean vs variance plot of mRNA expression values for 3’UTR-binding RNA-binding protein genes in adult mouse sensory neurons. X’s are individual genes. FXR1 and CELF4 are highlighted for reference. **B** Expression of a representative RBP, FXR1 is broadly expressed in all clusters of DRG neurons. Yellow circles are individual neurons that express FXR1 to high levels, blue circles = cells with low FXR1 expression **C** Heatmap of expression values of 3’UTR-binding RBPs significantly differentially expressed (FDR<0.01, top-10) in distinct populations of sensory neurons. Each line shows expression values in a single cell. Yellow = high expression, Purple = low expression. **D** Expression of significantly differentially expressed CELF-family RBPs in DRG neuron clusters. Yellow = high expression, blue = low expression. **E** Violin plots showing the distribution of expression values for significantly differentially expressed CELF family proteins. Yellow triangles denote significant enrichment, purple triangles denote significant exclusion. TH = tyrosine hydroxylase–expressing, NP = non-peptidergic, NF = neurofilament heavy chain-expressing and PEP = peptidergic neurons.

During these studies, numerous single-cell/nucleus sequencing studies were published and the data made publicly available. We therefore sought to confirm our findings using single nucleus sequencing data from adult mouse DRG (Renthal et al., 2020), and adult macaque (Kupari et al., 2021). We observed that CELF4 is predominantly in peptidergic populations (Fig. 3 a,b) confirming our previous results. Interestingly, we also observed that CELF1 and 2 are expressed in satellite/glial cells in addition to neurons (Figure 3 a).

**Figure 3:**
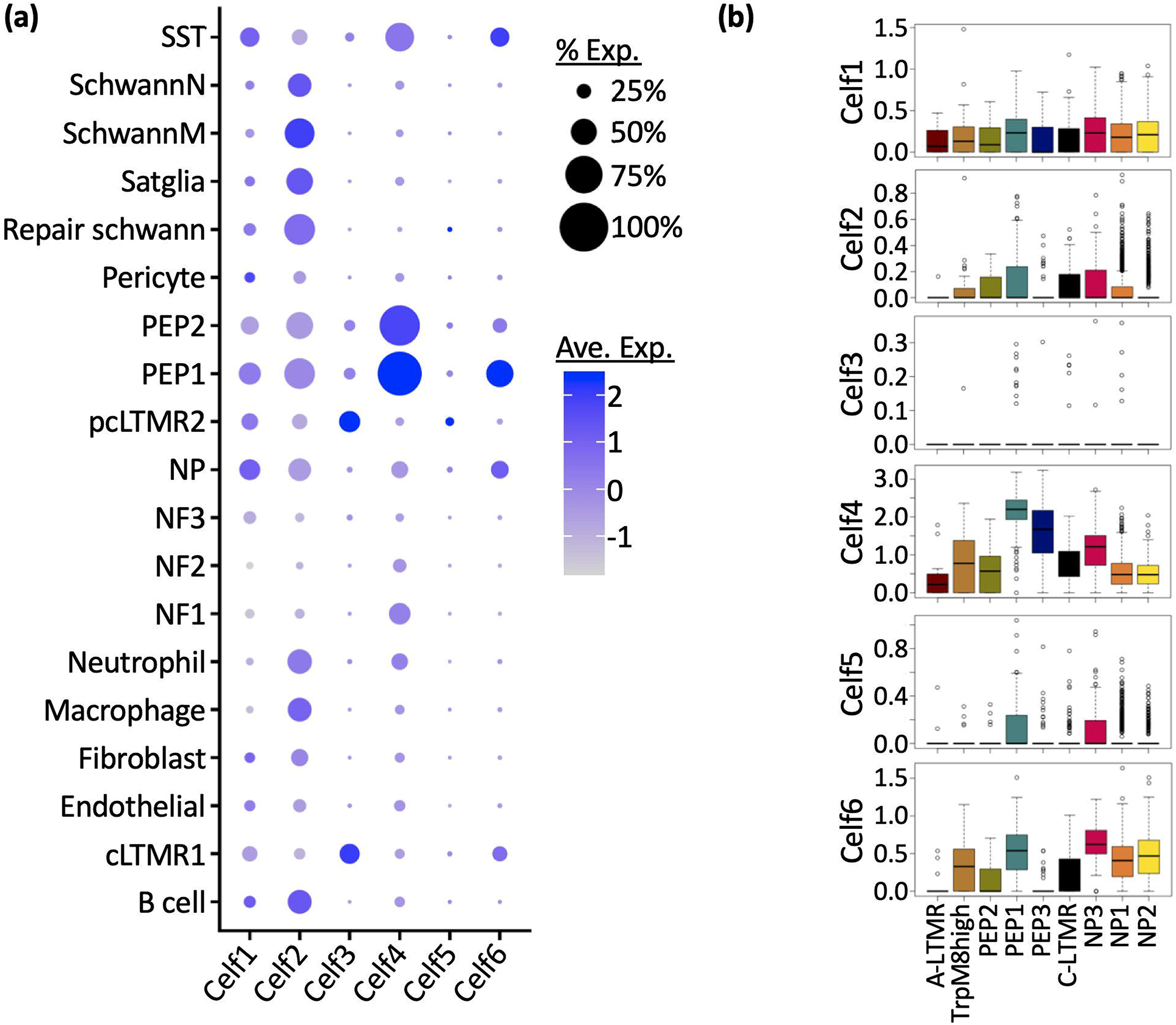
Re-analysis of publicly available single cell RNA-sequencing datasets confirm that CELF4 members are expressed in distinct populations of cells in sensory ganglia of adult mouse and macaque. **A** Expression of CELF member mRNAs in populations of neuronal and non-neuronal adult mouse sensory ganglia cells. Data from GSE154659 (Renthal et al., 2020). CELF1 and 2 mRNAs are detected in non-neuronal populations, whereas CELF4 is enriched in peptidergic neurons. **B** In the adult macaque, neuronal expression of CELF members is highly similar to mouse. Data from (Kupari et al., 2021).

CELF4 had the highest mean expression value of all family members (Figs. 2,3) and the largest differential expression value (28.2 fold) over the mean (Table 3). This family member negatively regulates neuron excitability in excitatory CNS neurons, likely by limiting translation of mRNAs encoding synaptic proteins and ion channels (Wagnon et al., 2011, 2012). However, expression and function of CELF4 in sensory neurons has not been described. RNA concentration does not necessarily correlate with protein concentration, e.g.:(Edfors et al., 2016). Therefore, the expression of CELF4 protein was examined using quantitative immunofluorescence microscopy. By measurement of fluorescent signals from individual somata from 3 animals (approx. 200 somata/animal), we observed that 58% of small diameter (<300 μm) and 38% of medium diameter (300-700 μm) DRG neurons were CELF4 positive (Fig 4). Using established histological markers for non-peptidergic (plant lectin IB4), peptidergic (CGRP antiserum) and myelinated (NFH antiserum) neurons, we observed that CELF4 protein distribution mirrors that of CELF4 mRNA. CELF4 was detected in 53% of CGRP+, 34% of IB4-binding and 54% NF-H+ neurons (Fig 5). However, CELF4 protein is expressed to significantly lower levels in IB4-binding (p<0.05, n=3) and NF-H+ (p<0.01, n=3 mice) neurons.

**Figure 4:**
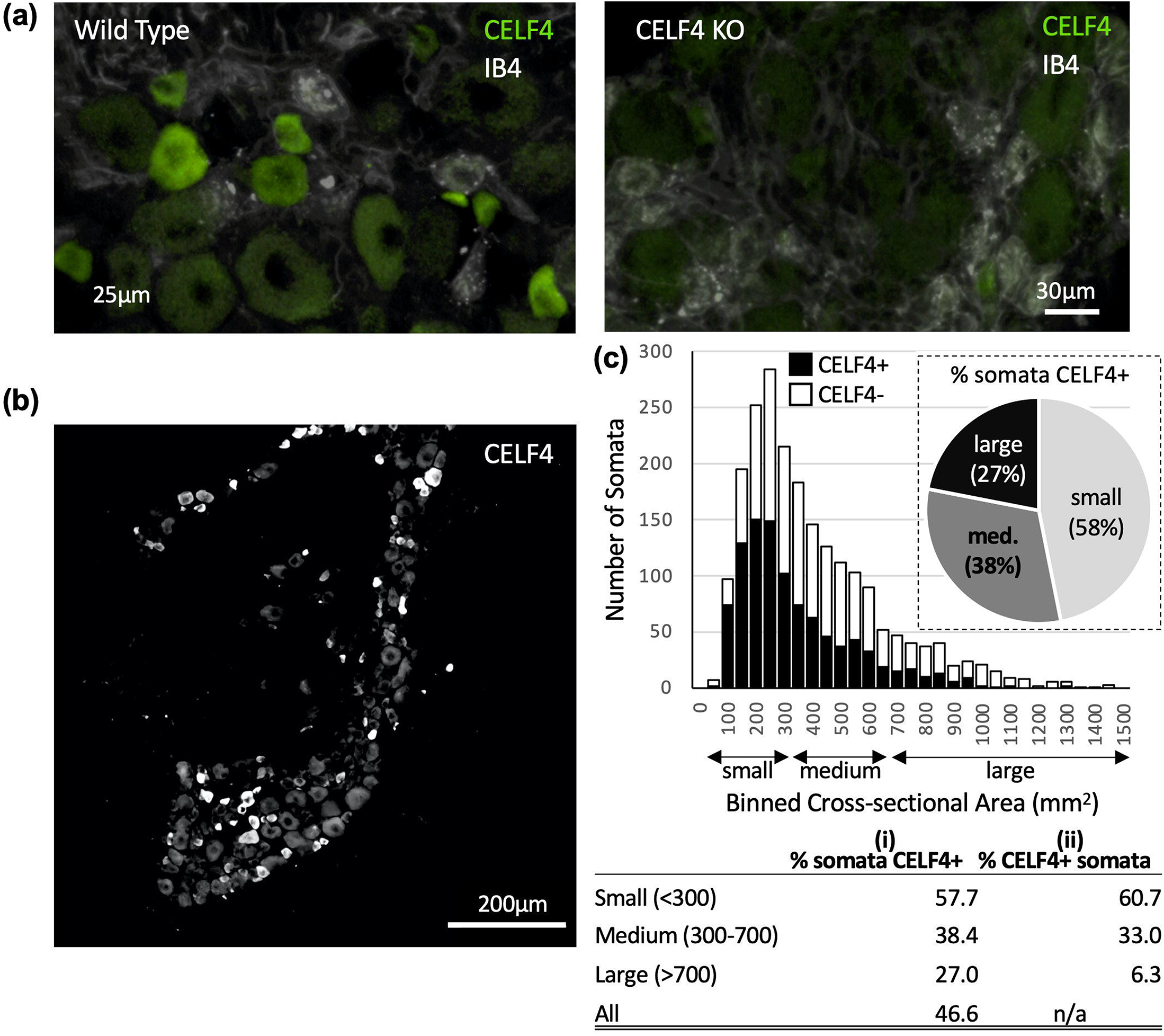
CELF4 protein is expressed in small and medium diameter sensory neurons. **A** CELF4 gene deletion abolishes CELF4 immunoreactivity, confirming the specificity of the CELF4 antibody. **B** Representative immunofluorescent micrograph of a 14μm DRG section stained with CELF4 antibody. **C** Soma size distribution of CELF4+ neurons. Data are collated from 3 sections, from 3 DRG from 3 replicate animals (approx. 100 soma per section). Somata were categorized according to cross-sectional area - small (<300μm), medium (300-700μm) or large (>700μm). **C i** Proportion of small, medium or large cross-sectional area DRG neurons that express CELF4. **ii** Proportion of CELF4-expressing neurons with small, medium or large cross-sectional areas.

**Figure 5:**
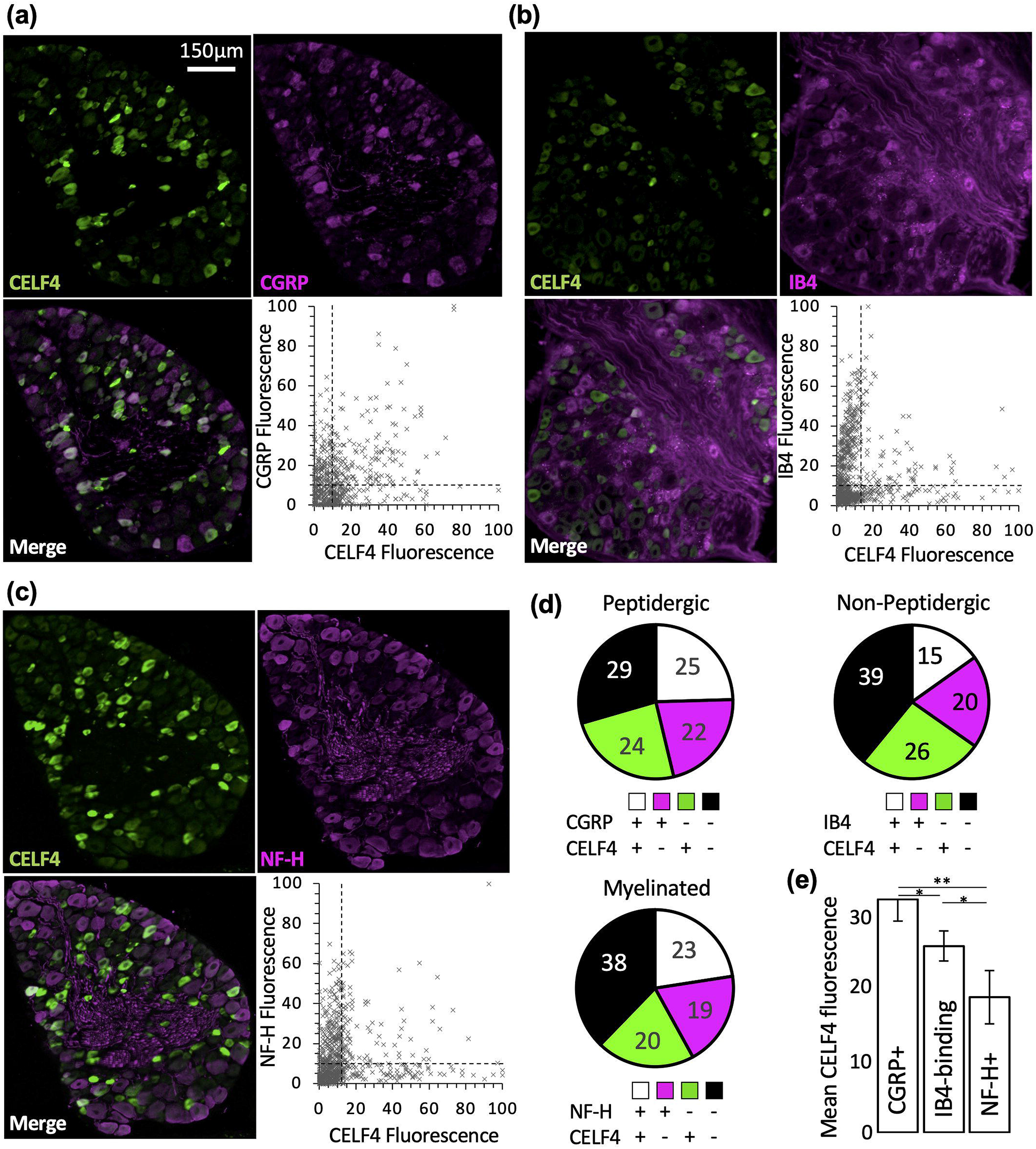
CELF4 protein expression in peptidergic (CGRP+), non-peptidergic (IB4-binding) and myelinated (NF-H+) neurons. **A-C** 14μm DRG sections stained with CELF4 antibody, co-stained with established histological markers. Peptidergic neurons were identified with CGRP antibody, non-peptidergic with isolectin IB4 and myelinated with NFH. Scatter plots show distribution of fluorescence intensities of somata collated from DRG from 3 replicate animals, 2 non-overlapping cross sections per animal (approx. 100 soma per section). Dashed lines indicate background signals levels. **D** Pie charts showing the proportion of double positive somata from each of the 3 stains. **E** Mean CELF4 fluorescence values. Stats = ANOVA with post hoc t-tests, n=3 animals, *=p<.05, **=p<.01.

By comparing CELF4 mRNA expression to that of all other mRNAs detected in the scRNA-Seq dataset, we determined that CELF4 expression is most highly correlated with GFRA3, TRPV1 and Calca (gene name for CGRP) expression in adult mouse DRG neurons (Fig 6, a).

**Figure 6:**
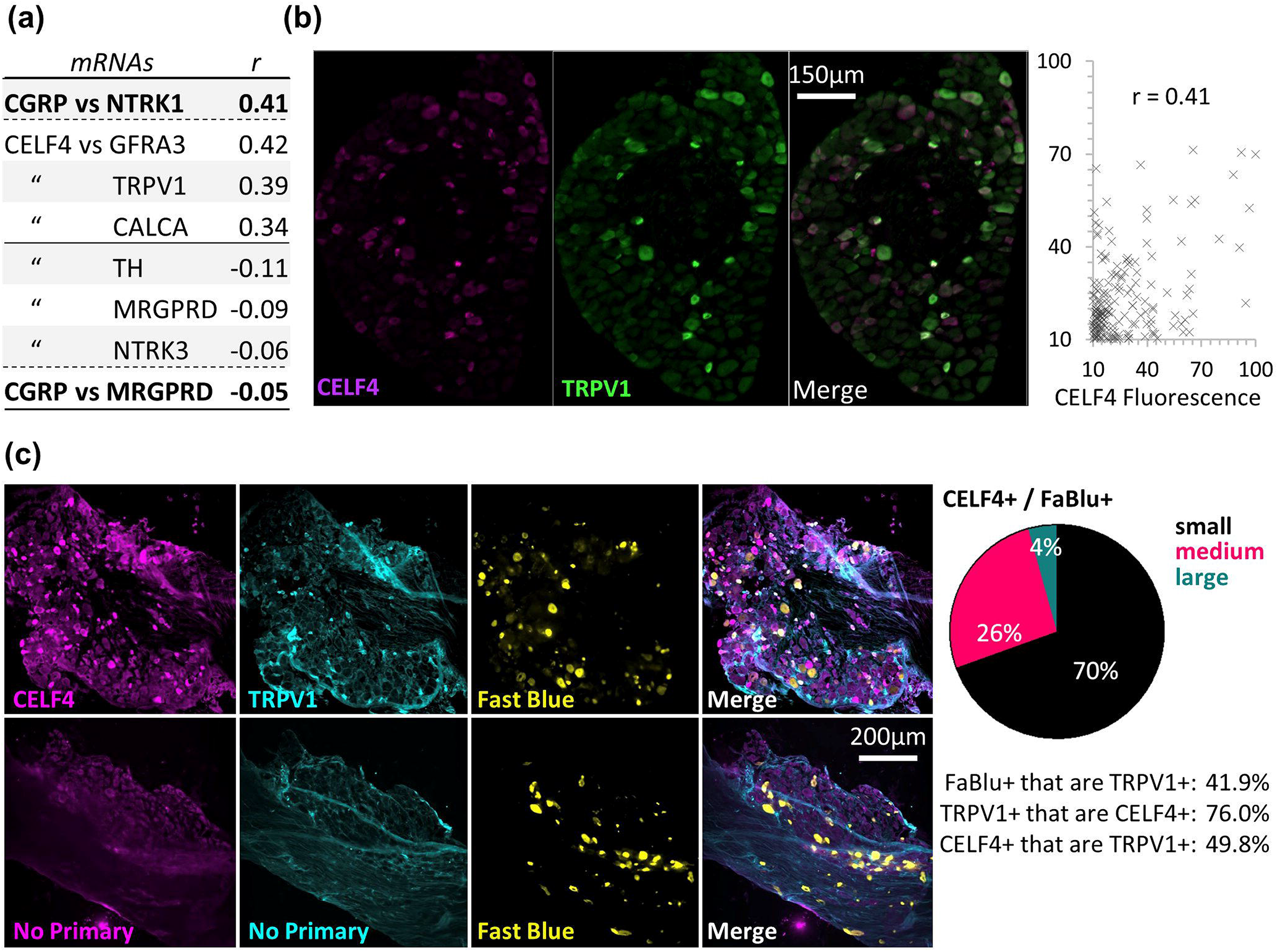
CELF4 is co-expressed with the TRPV1 capsaicin receptor in DRG neurons. **(A)** Pearson’s r correlation coefficients were calculated for CELF4 mRNA concentration vs all other transcripts. The 3 most positively and negatively correlated genes are shown. Also shown are reference values for a known co-expressed pair of mRNAs (CGRP vs NTRK1) and a pair of mRNA’s know to be expressed in different neurons (CGRP vs MRGPRD). (**B)** 14μm DRG sections were stained with CELF4 and TRPV1 antibody, and fluorescence intensity measured in soma of 2 sections from 3 animals. Representative micrographs are shown. (**C)** To support the mRNA correlations, we identified muscle afferents that co-express TRPV1 and CELF4 protein by using retrograde neuronal tracing from the gastrocnemius and tibialis anterior muscles combined with immunohistochemistry. Data collected from 3 DRG from 8 replicate animals (lumbar level 4). Representative micrographs are shown along with the no primary controls.

Given the prominent functional role of TRPV1+ afferents in muscle nociception and mechanical hyperalgesia, we chose to further characterize CELF4 protein expression in peripheral targets by taking advantage of the well-known central and peripheral innervation pattern of the sciatic nerve. Lumbar DRGs 3-5 contribute sensory innervation to peripheral targets to the muscles, vasculature, and skin of the leg. We first performed retrograde tracing from the tibialis anterior and gastrocnemius muscles to specifically label the DRG afferents innervating these muscles. In total, we characterized 2,490 (1 DRG each from 8 mice) retrogradely labeled primary sensory neurons for their expression of CELF4 and TRPV1. Many DRG neurons were CELF4+ (n=4876), and approximately 30% of them were also retrogradely labeled. We found that 42% of retrogradely labeled afferents also expressed the neuropeptide TRPV1 in L4 DRG, which agrees with previous findings using femoral nerve tracing and L2 DRG (Christianson et al., 2006). The majority of triple labeled (fast blue, CELF4, TRPV1) neurons was small in diameter. By co-staining DRG sections with TRPV1 and CELF4 antisera, we confirmed that the majority CELF4 immunofluorescence signal intensity is correlated with TRPV1 signal intensity (Fig 6b, c). Furthermore, among all TRPV1+ muscle afferents in the L4 DRG, 76% were also CELF4+.

Again, using our predicted CELF4 phenotype (primarily peptidergic – GFRA3, CGRP, TRPV1), we investigated CELF4 expression in another peripheral sensory target. Lanceolate endings are innervated by low threshold mechanoreceptors (LTMRs) whose cell bodies reside in the DRG. These specialized sensory termini are multiply innervated by a combination of Aβ−, Aδ−, and C-LTMRs. The specialized combinations of these sensory endings encode distinct sensory properties based on their myelination status and adaptation rates to sustained stimuli with at least 5 physiologically distinct LTMRs in hairy skin (Kuehn et al., 2019). Given that 90% of the body is covered with hair in most mammals, lanceolate endings are an indispensable unit for the perception of mechanosensory stimuli upon the skin. The majority of these lanceolate endings are also innervated by CGRP+ peptidergic nociceptors, which respond rapidly to hair pull and are developmentally dependent on NGF-TRKA signaling (Ghitani et al., 2017; Li & Ginty, 2014). Together with our finding that CELF4 is expressed in the cell bodies of this population, and that CELF4 is transported to neuropil in the CNS (Wagnon et al., 2012), we reasoned that a large proportion of lanceolate sensory endings in hairy skin would also express CELF4. We identified 114 lanceolate endings ~equally sampled among 5 mice, and 49.2% expressed CELF4. The pattern of CELF4 labeling suggests that CELF4 is present in both the circumferential and longitudinal endings surrounding the hair follicle (Fig 7), suggesting that CELF4 is present in the CGRP+ endings and/or at least one population of LTRMs

**Figure 7:**
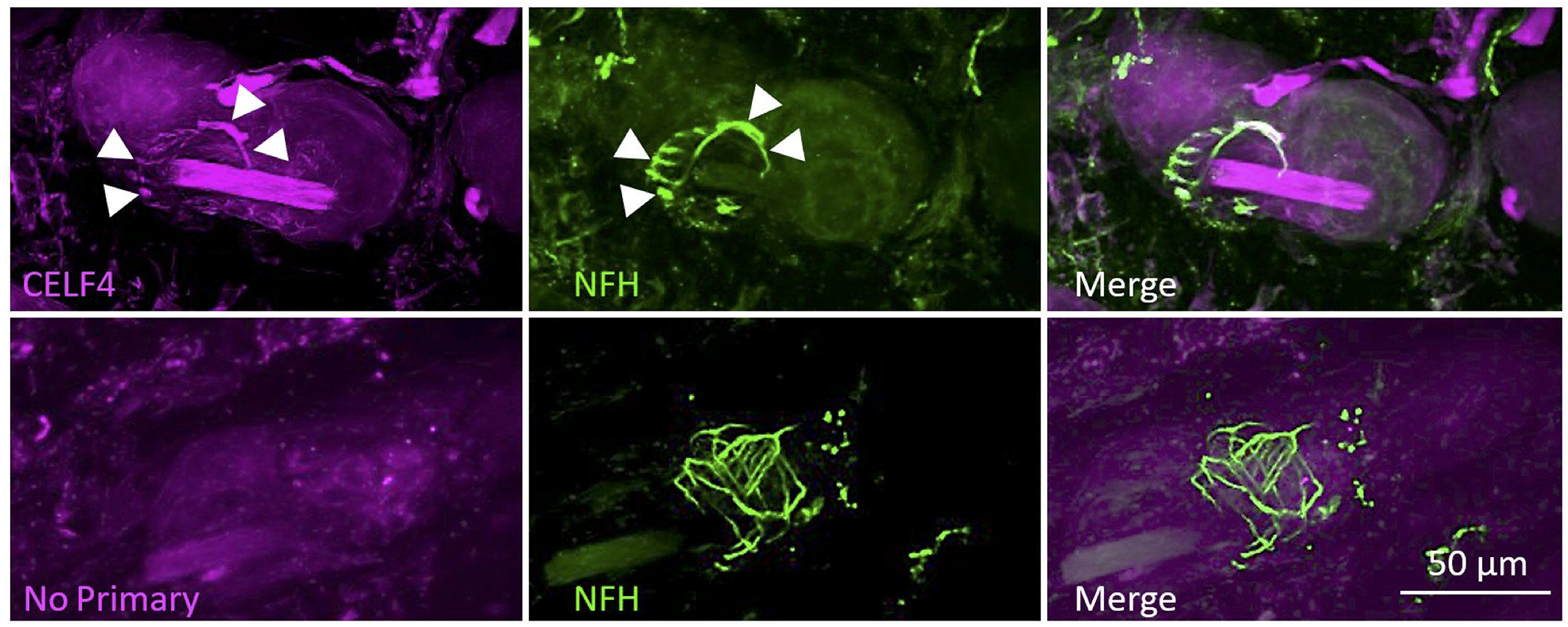
CELF4 is distributed to the peripheral termini of sensory neurons. The location of lanceolate endings in mouse hind paw hairy skin sections was identified by NFH immunoreactivity. CELF4 was detected in 49.2% of lanceolate sensory nerve endings in hair follicles within both the longitudinal and circumferential portions of the sensory terminals (arrow heads). Representative micrographs are shown along with no primary control. n=5 animals.

Previous work has shown that CELF4 regulates neuron excitability (Wagnon et al., 2011) and that CELF members may play a role in axon growth (Chen et al., 2016). We therefore wanted to examine the expression of CELF proteins in a model of nerve injury where hyperexcitable neurons are regenerating. Re-analysis of data from sciatic crush (Renthal et al., 2020) strongly indicated that CELF RNA-binding proteins are differentially expressed during the time course of recovery from sciatic injury (Fig. 8a). This was confirmed by analysis of bulk RNA-seq data derived from rat sensory ganglia post sciatic injury (Harrison et al., 2019) where CELFs are down-regulated during the course of recovery (Fig. 8 b).

**Figure 8:**
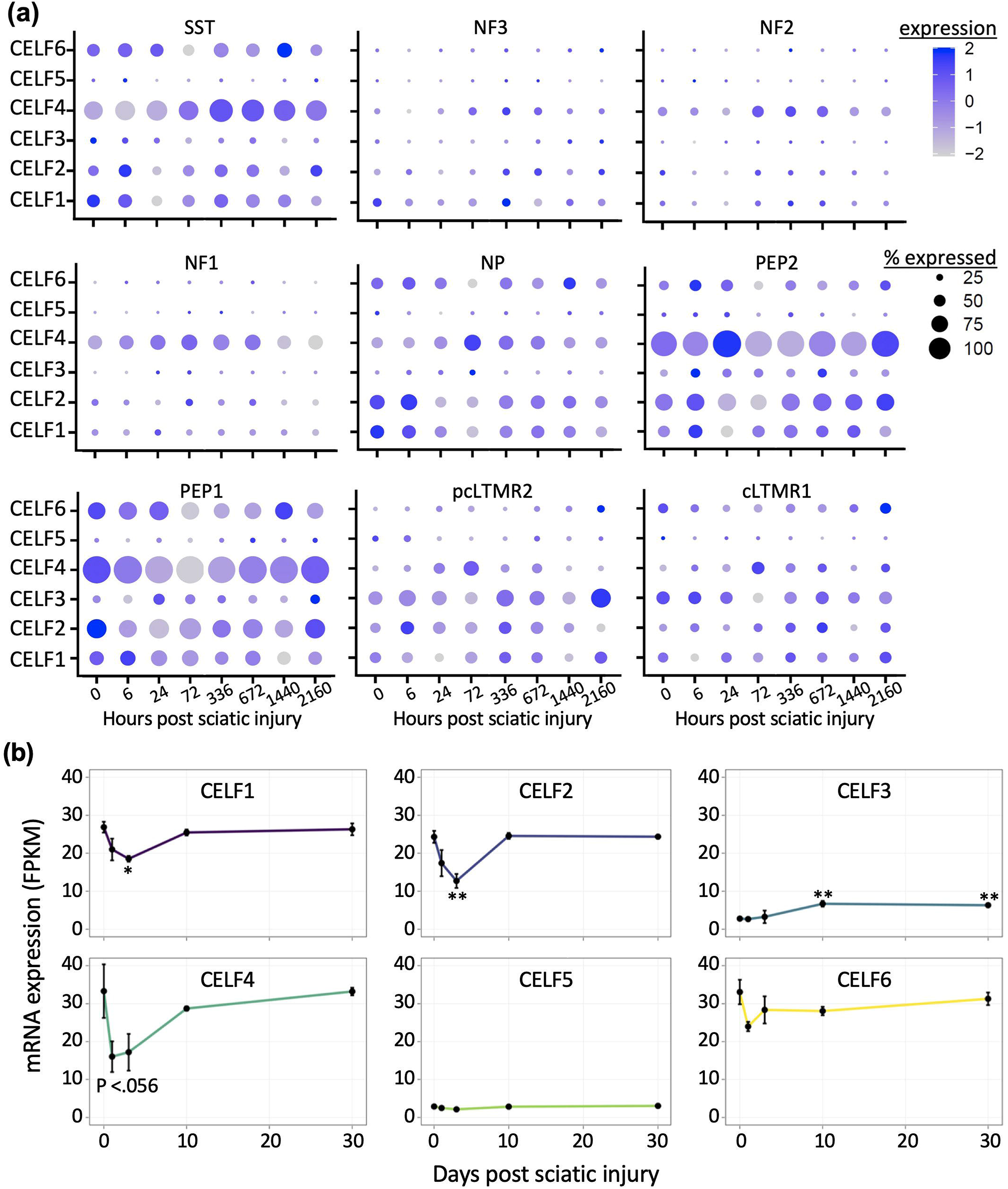
CELF transcripts are downregulated in sensory ganglia in rodent models of neuropathic pain. Single nucleus sequencing data GSE154659 (Renthal et al., 2020) from mouse DRG (**A**) and bulk homogenate RNA-Seq from rat DRG (Harrison et al., 2019) (**B**) showed differential expression of CELF members post sciatic injury. * = q<0.05, ** = q<0.01 where q values are determined by Student’s t-tests with Bonferroni multiple testing correction.

## DISCUSSION

Our previous work has shown that populations of sensory neurons express different mRNA isoforms with alternate 3’UTRs (Harrison et al., 2014). In the current study, we investigated the expression of 3’UTR-binding proteins in DRG, leading to the discovery that members of the CUGBP-binding ELAV-like family of RBPs are expressed in distinct populations of sensory neurons. By analysis of publicly available scRNA-Seq data, we revealed that CELF2 is expressed in small diameter neurons and non-neuronal cells, CELF4 is restricted to the peptidergic (PEP) population, and CELF6 is present in both peptidergic and non-peptidergic (NP) populations. These neuronal populations predominantly represent different classes of C-fiber nociceptors, and in the case of PEP also a subset of A-delta fibers. Most CGRP-expressing (PEP) C-fibers also express the pro-inflammatory neuropeptide substance P (Juránek & Lembeck, 1997). CGRP and SP are pro-inflammatory peptides that play key roles in neurogenic inflammation; CGRP is a potent vasodilator and substance P induces mast cell degranulation (Lim et al., 2017). In contrast, NP neurons have been reported to suppress mast cell reactivity (Zhang et al., 2021). Notably, several studies have demonstrated that cutaneous IB4-binding neurons transduce mechanical stimuli in the non-noxious range, and optogenetic stimulation of these neurons fails to evoke nocifensive behavior unless they are sensitized by tissue injury, calling into question whether these neurons act as nociceptors in healthy tissue (Warwick et al., 2021). CELF3 is restricted to the TH population of non-peptidergic, innocuous low threshold C-fiber mechanoreceptors, which have been implicated in the pleasurable, affective component of touch (Lallemend & Ernfors, 2012; Li et al., 2011). Immunostaining revealed that the family member with the highest expression, CELF4 is enriched in neurons that express the capsaicin receptor TRPV1 and are therefore responsive to noxious heat. CELF4 is a negative regulator of mRNA translation and neural excitability (Wagnon et al., 2011). We therefore propose that further study of CELF4 could uncover mechanisms that regulate protein synthesis in sensory neurons that convey pain and are important for neurogenic inflammation.

Previous studies examining the expression profile and function of CELF members have determined that these proteins are expressed in the central nervous system during development and in the adult (Table 4) (Giudice et al., 2016; Krismer et al., 2020; Samaras et al., 2020; Schmidt et al., 2018). The RNA-binding motifs for all members are similar, with UG repeats (Table 4), suggesting that these proteins have the ability to bind similar transcripts. However, given that expression of CELF members is restricted in a cell-type dependent manner, this could limit functional redundancy between members. Focusing on CELF4, in the CNS this member is expressed primarily in excitatory neurons, including large pyramidal cells of the cerebral cortex and hippocampus, where it tonically limits excitatory neurotransmission (Wagnon et al., 2012). In these neurons, RNA-protein interaction profiling (RIP-Seq) revealed that CELF4 predominantly binds to the 3’UTR of transcripts, and CELF4 knockout reduced mRNA stability (Wagnon et al., 2012), and resulted in increases in expression of sodium channel Na(v)1.6 (Sun et al., 2013). Therefore, CELF4 is a negative regulator of excitability of excitatory CNS neurons most likely though limiting translation in these neurons. In accordance, electrophysiological experiments using brain slices revealed that loss of CELF4 function lowered the action potential (AP) initiation threshold and increased AP gain (Sun et al., 2013).

**Table 4:** CELF family RNA-binding protein tissue distribution, subcellular location, RNA recognition motifs and summarized function. Tissue expression data from ProteomicsDB <u>(Samaras et al., 2020; Schmidt et al., 2018)</u>. RNA-binding motifs were obtained from Transite <u>(Krismer et al., 2020)</u> and ATtRACT <u>(Giudice et al., 2016)</u> databases. * reported consensus sequence

| *Family member* | *Tissue Expression (Adult)* | *Subcellular Location* | *RNA-binding motif* | *Molecular/biological function(s)* |
| --- | --- | --- | --- | --- |
| CELF1 | Brain (m), spleen (m), liver(m), testis(m) ubiquitous(h) | Nuclear and cytoplasmic | UGUGUG,UGUU,<br>CUGUCUG,UUGUG, | splicing,RNA stability, translation regulation (1-5) |
| CELF2 | Brain (m), spleen (m),heart(m), lung(m),testis(m), ubiquitous(h) | Nucleoplasm, vesicles | UGUGU, UUGUU,UGUU, | RNA binding, mRNA processing,splicing |
| CELF3 | Brain (m), adrenal gland (h),retina (h), pancreas (h) | Nucleus, cytoplasm | UGUUGUG (h) | RNA binding,mRNA splicing,spermatogenesis |
| CELF4 | Brain (m,h), adrenal gland (h), retina (h) | Nucleus, cytoplasm, Axons/Dendrites | UGUGUKK(h), UGUGUGU* | RNA binding, mRNA processing, mRNA splicing |
| CELF5 | Brain (m), N/A(h) | Nucleus, cytoplasm | UGUGUKK (h), UGUGUGU* | RNA binding |
| CELF6 | Brain (m), N/A (h) | Nucleus, cytoplasm, synapse | UGUGKKG (h) | RNA binding, mRNA processing |

This is the first report characterizing expression of CELF4 in populations of neurons in the peripheral nervous system, and its function in those neurons remains unknown. Extrapolating from studies performed with CNS neurons where CELF4 primarily binds to sites within the 3’UTR of mRNAs, indicated molecular functions of this protein could include alternate polyadenylation thereby coordinating the expression of 3’UTR isoforms. This would be an enticing possibility due to our previous observations of differential 3’UTR isoform expression in DRG neuron populations (Harrison et al., 2014), and mechanisms responsible for cell-type specific expression of 3’UTR variants are currently unknown. However, in the current study we did not detect CELF4 in the nucleus of DRG neurons, making this possibility less likely. Also, in the CNS CELF4 mRNA co-sediments with polysomes purified from neuropil, and CELF4 knockout increases the concentration of key excitability proteins in this compartment (Wagnon et al., 2012). Therefore, CELF4 limits local translation in CNS axons. Together with our findings, this suggests that CELF4 could also limit protein expression in populations of sensory neuron termini, and experiments to characterize CELF4 expression in central and peripheral endings are currently underway.

CELF4 tonically limits protein translation in and excitability of neurons. Therefore, pharmacological targeting of CELF4-3’UTR interactions could lead to the development of tools to study mechanisms of protein translation in neurons. Targeting CELF4 could have wide-ranging implications for diseases of the nervous system. For example in the CNS, CELF4 knockout mice develop a seizure phenotype closely resembling an epilepsy syndrome caused by CELF4 locus (18q12) deletion in humans (Halgren et al., 2012). Therefore, enhancing CELF4 function has been proposed as a strategy to treat epilepsy. We have determined that in the PNS CELF4 is enriched in C/Aδ neurons, a population of neurons critical for the etiology of pathological pain, and that this protein is regulated during the time course of recovery from sciatic injury. Therefore, enhancing CELF4 function in nociceptors could be a novel approach to preventing/alleviating chronic pain.

## ACKNOWLEDGMENTS

P20GM103643 MH116492 T32GM107000

